# Phylogenetics, Epidemiology and Temporal Patterns of Dengue Virus type 1 in Araraquara, São Paulo State

**DOI:** 10.1101/2023.09.18.558374

**Authors:** Caio dos Santos Souza, Giovana Santos Caleiro, Ingra Morales Claro, Jaqueline Goes de Jesus, Thaís Moura Coletti, Camila Alves Maia da Silva, Angela Aparecida Costa, Marta Inenami, Andreia C Ribeiro, Alvina Clara Felix, Anderson Vicente de Paula, Walter M Figueiredo, Expedito José de Albuquerque Luna, Ester Sabino, Camila Malta Romano

## Abstract

Dengue virus is a prominent arbovirus with global spread, causing approximately 390 million infections each year. In Brazil, yearly epidemics follow a well-documented pattern of serotype replacement every three to four years on average. Araraquara, located in the state of São Paulo, has faced significant impacts from DENV epidemics since the emergence of DENV-1 in 2010. The municipality then transitioned from low to moderate endemicity in less than 10 years. Yet, there remains an insufficient understanding of the virus circulation dynamics, particularly concerning DENV-1, in the region, as well as the genetic characteristics of the virus. To address this, we generated 37 complete or partial DENV-1 genomes sampled from 2015 to 2022 in Araraquara. Then, using also Brazilian and worldwide DENV-1 sequences we reconstructed the evolutionary history of DENV-1 in Araraquara and estimated the time to the most recent common ancestor (tMRCA) for the serotype 1, for the genotype V and its main lineages. Within the last ten years, there were at least three introductions of genotype V in Araraquara, distributed in the two main lineages (LIa and LIb, and LII). The tMRCA for the first sampled lineage (2015/2016 epidemics) was approximately 15 years ago (in 2008). Crucially, our analysis challenges existing assumptions regarding emergence time of the DENV-1 genotypes, suggesting that genotype V might have diverged more recently than previously described. The presence of the two lineages of genotype V in the municipality might have contributed to the extended persistence of DENV-1 in the region.

## 1.0 Introduction

According to the World Health Organization (WHO), Dengue Virus (DENV) stands as the most relevant arbovirus in the world, posing a threat to 3,9 billion people in 120 countries [1]. Annually, over 390 million infections are estimated to occur, with tropical and subtropical areas being the most affected ones, particularly in urban and developing regions such as countries in South America and Asia [1].

DENV belongs to the *Flavivirus* genus and *Flaviviridae* family and is classified in 4 serotypes (DENV-1 to DENV-4) due to antigenic and genetic differences, with each serotype further exhibiting various genotypes [2]. Infection with any serotype can lead to dengue fever, a disease whose clinical manifestation ranges from flu-like symptoms to a life-threatening condition known as severe dengue [1]. In Brazil, from 2013 to 2022, a substantial number of possible dengue cases amounting to 9,938,730, with 5,836 deaths were reported [3,4]. During this period, there was a clear pattern of alternating circulating serotypes between DENV-1 and DENV-2 in the country, except for the year 2013, when DENV-4 was the main circulating serotype [3,4]. In this scenario, the municipality of Araraquara emerged as a significant contributor to the escalating number of dengue cases [5].

Up until 2005, only sporadic cases were reported in the municipality. However, the panorama of dengue fever underwent a significant change in 2006, when serotype 3 was introduced into the region, leading to the first reported outbreak [5,6].

Subsequently, in 2010, the introduction of serotypes 1 and 2 resulted in a significant raise in the number of cases and deaths [5]. Particularly noteworthy were the years 2015, 2019 and 2022, which reported 8,209; 23,538; and 21,070 cases, respectively [7]. During the period of 2014-2018, DENV-1 was the main serotype circulating in the municipality, followed by clade replacement in 2019, when DENV-2 triggered a massive outbreak [8].

In this study, we performed genome sequencing analysis of DENV using samples collected from the municipality of Araraquara in 2015, 2016, 2019, 2021, and 2022 to assess the molecular characteristics of the circulating viral strains during this period.

## 2.0 Methods

### 2.1 Ethical statement

This study was approved by the ethical committee of the Medicine school of University of São Paulo, Approval under Nº 4,519,364, encompassing authorization for sample collection, viral detection and genomes sequencing.

### 2.2 Study population

The study participants are part of a cohort that has been under observation since 2014, originating from the municipality of Araraquara, São Paulo. The collection of samples followed specific criteria linked to the appearance of symptoms suggestive of dengue fever [8]. During the initial week following symptom onset, plasma and serum samples were gathered and subsequently forwarded to the virology laboratory at the Institute of Tropical Medicine (IMT-USP).

### 2.3 DENV RNA detection and sample selection for Next Generation Sequencing (NGS)

Viral RNA was extracted from 140μl of plasma and/or serum using the QIAamp Viral RNA MiniKit (QIAGEN, Hilden, Germany) according to the manufacturer’s instructions, obtaining a total of 60μl of RNA. For the initial detection of viral RNA, a probe-based qPCR capable of detecting the 4 DENV serotypes was performed [9,10]. Following the confirmation of DENV infection, samples were typed through a specific qPCR [10,11]. DENV-1 samples with the cycle threshold (CT) < 30 were eligible for the Next-Generation Sequencing (NGS).

### 2.4 DENV-1 complete genome amplification

The primers to amplify the complete genome of DENV-1 were designed using the online tool PrimalScheme (available at primalscheme.com) and a polifasta file containing reference DENV-1 complete genomes as input. After defining the desired amplicon size of 400 base pairs and an overlap of 75 nucleotides, two sets of primers suitable for implementation in a multiplex scheme were generated, arranged into two pools [12].

Following the cDNA synthesis utilizing First Strand cDNA ProtoScript II (New England Biolabs, Oxford, England), the two multiplex PCR reactions were conducted under the conditions described elsewhere [13]. Amplicons were purified using 1X AMPure beads (Beckman Coulter, United Kingdom) and quantified using the Qubit 4 fluorometer (Invitrogen, California, USA). DNA concentration >2ng DNA/μl per sample was considered suitable for whole genome sequencing.

The libraries were generated using the genomic DNA sequencing kit SQK-LSK108, where 250ng of Barcoded PCR products were pooled in equimolar proportions. The libraries were loaded onto an Oxford Nanopore Flow Cell R9.4 and the data were collected after ∼30h [13].

### 2.5 Bioinformatics workflow and consensus generation

The basecalling of the FAST5 reads containing the raw signal data was done using Guppy basecaller integrated within MinKNOW (https://community.nanoporetech.com/). Then the FASTQ files were used to map the reads using the genome ID#KP188547 as reference in the CLC genomics workbench (Qiagen, Hilden, Germany) [14]. Genomes with coverage >10 per base were exported as FASTA format for further analyses.

### 2.6 Envelope gene sequencing

The DENV genomes that were not suitable for Next-Generation Sequencing (NGS) underwent partial envelope sequencing using the Sanger method. We decided to do this to include the maximum number of samples from Araraquara in the analysis. To amplify ∼800pb of the envelope, we performed a PCR using primers and protocol described elsewhere with modifications [15]. Briefly, the PCR reaction was performed using 14.75μl of Ultrapure Nuclease Free Water (NFW), 1x PCR Buffer, MgCl^2^ 1.5mM, 0.25uM of each primer, 0.4μM of dNTP and 1U of PlatinumTaq Polymerase in a final volume of 25μl.

Amplicons were sequenced using BigDye Terminator v3.1 on the ABI 3730 DNA analyzer (Thermo Fisher Scientific, Massachusetts, USA). Electropherograms were analyzed and the consensus sequences were assembled using the Geneious Prime software (Dotmatics, Hertfordshire, England).

### 2.7 Multiple sequence alignment and ML Phylogenetic reconstruction

The dataset comprised 284 partial E gene sequences (∼800 nucleotides long) retrieved from GenBank (Sequence list available at supplementary material-Table S1) containing sampling date, host and country information and 37 sequences generated in this study (either by NGS or Sanger). Sequences were multiple aligned using MAFFT v7.407 and visually inspected using the SeaView v5.0.4 software [16,17]. To first classify DENV-1 sequences into genotypes we reconstructed a Maximum Likelihood phylogenetic tree using the IQ-TREE 2 under the GTR + I + G nucleotide substitution model, as the best-fitting one selected by ModelFinder implemented in IQTREE [18]. The phylogenetic tree was visualized with FigTree v.1.4.4 [19].

### 2.8 Coalescent analysis and molecular clock

The time to most recent common ancestor (tMRCA) of the DENV-1 genotypes, and particularly the Araraquara ones were estimated using the Bayesian Markov Chain Monte Carlo implemented in BEAST 1.10.4 [20,21]. The Bayesian Skyline (BSL) coalescent method was performed under relaxed uncorrelated exponential molecular clock using time-stamped data scaled in years under GTR + I + G nucleotide substitution model. Convergence of parameters was inspected with Tracer v.1.7.2 with uncertainties addressed as 95% highest probability density (HPD) intervals.

After 20 million runs, the trees sampled at every 2,000 steps were summarized in a maximum clade credibility (MCC) tree with 10% burn-in using TreeAnnotator and visualized with FigTree v.1.4.4. In addition to the BSL method, the exponential growth tree prior was run for comparison.

## 3.0 Results

A total of 37 samples were positive for DENV-1 with cT < 30, median 24.5 (18.4 - 30.2) and were submitted to the whole genome amplification. The number of samples per year was: 2015 (N=26), 2016 (N=1), 2019 (N=2), 2021 (N=2) and 2022 (N=6). All samples were collected during the first semester of the specific years.

Twenty-eight out of the 37 samples submitted to the whole genome amplification had enough DNA concentration and were sequenced using Nanopore method. The remaining samples (N=9) were submitted to Sanger sequencing (envelope gene). The sequences were submitted to GenBank database under accession numbers: OR553381-OR553391; OR348411-OR348418; OR545518-OR545534; and OR520604.

Maximum likelihood analysis of DENV-1 showed a clear separation of the distinct genotypes and the genotype V division in two previously classified lineage I and lineage II described by Bruycker-Nogueira and colleagues [22] (data not shown). We then run the same dataset in BEAST to perform the temporal estimates. According to the BLS estimates, DENV-1 coalesced approximately 139 years ago (95% HPD 103 -190 ya), in 1884, while the origin of the Genotype V was estimated to be around 58 years ago (95% HPD 49 – 62 ya), in 1965. Similar data was found under the exponential growth. All estimates with 95% HPD for both coalescent models are presented in Table 1.

The Maximum Clade credibility (MCC) tree also showed the 2 different lineages of genotype V [22]. The genomes from 2015 and 2016 were classified as lineage II (LII), while genomes collected from 2019 onwards were classified as lineage I (LI). According to Bayesian estimates, the Lineage II coalesced in 1982 (around 41 years ago); Lineage I coalesced in early 1971 (around 52 years ago), and lineages Ia and Ib coalesced in 2001 and 1996 respectively (Figure 1 and Table 1).

**Figure 1:**
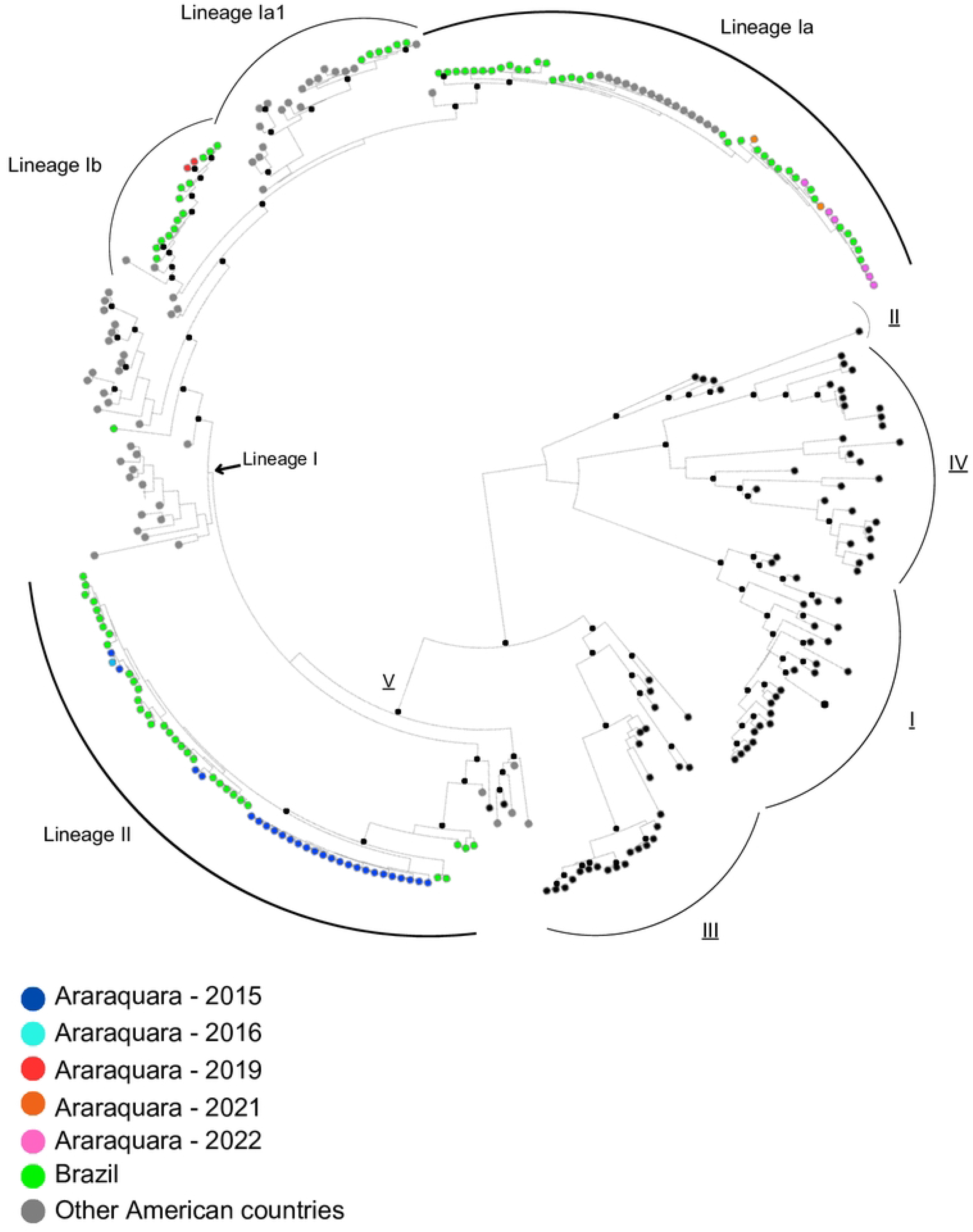
Maximum clade credibility tree of DENV-1. The genotypes I to IV (black circles) are identified by underlined roman numerals. The genotype V that includes most of the reference sequences and all Brazilian DENV-1 is also classified in lineages and sub-lineages. Small black dots denote the nodes with posterior probability > 0.7. The taxa were colored according to the legend.

Supported by high posterior probability values, we identified an intermediate clade between the previously classified lineages Ia and Ib, named here as Ia.1. This clade comprises viruses from South and Central America and Caribbean, being the most ancient virus sampled in Colombia, in 2014. Brazilian sequences within clade Ia.1 are more recent, and were all sampled in São Paulo, from 2021 to 2023. According to our estimates, the clade Ia.1 emerged late in 1997.

All but 5 viruses from Araraquara 2015/2016 clustered together in a branch that probably emerged in 2008 (95% HPD 2005-2011). Two out of the remaining samples from 2015 clustered with samples from Goiás (2013) and Rio de Janeiro (2013). This second cluster consists primarily of Brazilian DENV strains, comprising viruses from Goiás and Rio de Janeiro (2013), Ceará and Sergipe (2014), and São Paulo (2016). Notably, these two clusters constitute the entire Lineage II.

Viruses from Araraquara 2019 belong to Lineage Ib and grouped to others viruses from São Paulo (2016 and 2018). The cluster also included viruses from the states of Roraima (2010), and Ceará (2014). The tMRCA for this group was estimated 5.5 years ago (95% HPD 4-7ya), in 2017. It is also worth noting that, in contrast to the lineage II, some of these genomes were closer to other South American countries, such as Venezuela and Colombia.

The DENV-1 sequences obtained in 2021 and 2022 clustered together and were close to other São Paulo (2021, 2022 and 2023), Rio de Janeiro (2016) and Mato Grosso do Sul (2016) strains. The node comprising these sequences is dated to 9.3 years ago (95% HPD 8-10 ya), in 2013.

Finally, the dates of key DENV-1 events were re-estimated. According to our results, DENV-1 had its origin 139 years ago, estimated in 1884 (95% HPD 1833-1920), and the genotype V arose 58 years ago, in 1965 (95% HPD 1955-1972). The approximated origin of the Genotype V lineages and sublineages as well as the tMRCA of the other four genotypes is described in the Table 1.

## 4.0 Discussion

The phylogenetic structure of dengue viruses in endemic regions is marked by lineage turnovers, when serotype or lineages are replaced every 3-4 years [3,23]. DENV serotype replacement has been long observed, and was reported in detail in Thailand, Malaysia, Vietnam, Panamá and other countries [23,24]. Brazil annually reports the highest number of dengue fever cases and deaths globally, and also presents a classical pattern of serotype and clade replacement over the years [3,25].

Besides serotype turnovers, replacement of genotypes and lineages is also common. This is often linked to higher case and death counts in endemic regions, likely resulting from varying susceptibilities to cross-reactive immune responses against the currently circulating virus [26,27].

Araraquara, located in the central region of São Paulo state is among the most industrialized cities in the state, and reported the first autochthonous dengue case in 1996. However, information regarding the serotype was not available until 2006, when DENV-3 was described in the region. Since then, the number of cases progressively rose leading to severe outbreaks and epidemics every year [5,7]. After the emergence of DENV-1 in 2010, Araraquara experienced an extended predominance of this serotype for several years with a remarkable number of reported cases over the years [7]. Particularly, from 2014 to 2018, DENV-1 emerged as the predominant serotype in Araraquara, while DENV-2 dominated from late 2018 and 2019, leading to more than 20 thousand reported cases [5,7,8]. Nonetheless, in 2020, DENV-1 reclaimed its status as the prevailing serotype and has continued to do so up to the present time [28].

The phylogenetic analysis using longitudinal DENV-1 samples collected in an 8 years time-span performed in this study represents the first description of DENV-1 lineages circulating in the municipality of Araraquara. During this period, we detected the presence of the two lineages of DENV-1 genotype V in Araraquara, L I and L II [22]. Initially, DENV-1 was represented exclusively by the lineage II and almost all clustered together. Only three samples (two from 2015 and one from 2016) did not cluster with the remaining viruses collected in 2015, though we cannot definitively determine whether this is due to a distinct introduction of a very similar strain or a result of locally accumulated diversity. From 2019 onwards, viruses from L II were no longer sampled, and all circulating viruses until 2022 belonged to the Lineage I. The phylogeny also evidenced a distinct cluster within Lineage I, sufficient divergent from L Ia and L Ib previously described [22], named here L Ia.1. Although this cluster comprises Brazilian viruses, no sample from Araraquara was classified as L Ia.1

Particularly, viruses from 2019 clustered within sub-clade L Ib, while those from 2021 and 2022 belonged to the sub-clade L Ia, revealing a second clade replacement. The presence of distinct lineages and sublineages of DENV-1 in Araraquara might have played a critical role in sustaining the prevalence of this serotype in the region for such an extended period, as it has been seen elsewhere [27]. Unfortunately, we don’t have samples prior to 2015 and also no samples were collected in 2017 and 2018, so it is impossible to precisely determine how many clade/lineages replacements occurred since DENV-1 emergence in Araraquara.

It has previously been proposed that homotypic lineages undergo antigenic evolution in such a way that they occasionally resemble lineages of different serotypes. This phenomenon is particularly marked for DENV-1 and DENV-2, which are also the most prevalent in Brazil [29]. It has also been demonstrated that certain homotypic lineages of DENV are not more effectively neutralized than heterotypic viruses [30] even after a two-year period, opening the possibility of homotypic reinfection if the second lineage is sufficiently different from the previous one. It would be the case for regions presenting continued circulation of distinct lineages from a specific serotype as Araraquara. Alternatively, at least for Araraquara, the demographic factor may also serve as a reasonable explanation, as when combining the number of potential cases during the two DENV-1 major epidemics (2015 and 2022), around one-fourth of the population would have been exposed to this serotype.

While several studies point that clade replacement is a stochastic event and does not depend on selection, there is also evidence that strains of DENV can exhibit different fitness either in mosquito or in vertebrate hosts [24,31,32,33]. For instance, during a clade replacement in Indonesia within the Cosmopolitan genotype of DENV-2, the increased number of cases was associated with a higher fitness of the new clade in mammalian cells [34].

Our group recently reported a clade replacement within DENV-2 American/Asian genotype (III) in São Paulo, where the introduction of the BR-4 lineage was associated with major outbreaks in 2019 and rapid expansion of the lineage in the country [26]. Although we have not evaluated the number of deaths and severe dengue cases in the 2015 and 2022 DENV-1 epidemics, it was noticeable the higher number of cases caused by the Lineage I compared to the Lineage II (i.e. 8,209 in 2015 and 21,070 reported cases in 2022).

Our bayesian analysis also enabled us to re-estimate the dates of key DENV-1 events. According to the estimates, the origin of the serotype 1 as well as the genotypes I to IV were similar to those reported elsewhere [36]. However, we noticed that the clustering pattern of two sequences (AF425620, from Cote d’Ivoire and AF425625, from Nigeria), previously classified as genotype V [36], were not consistent and grouped near genotype III in the ML (not showed) and MCC trees.

The previous reports that considered these sequences as genotype V dated the origin of this genotype back to 1935 [35,36]. However, according to our estimation, Genotype V might have emerged much more recently, around 1968. It has to be noticed that both previous studies included a limited number of Genotype III reference sequences. Consequently, the lower genetic diversity of Genotypes III and V from the previous phylogenetic analysis might have hindered accurate classification during the reconstructions.

In an effort to explore the potential factors underlying the discrepancies observed in previous phylogenetic classifications, we tested these samples for recombination, and estimated the genetic distances between them and others from Genotypes III and V. RDP [37] found no evidence of recombination in both sequences. Genetic distance analysis using the Kimura 2-parameter revealed >6% distance between these genomes and those from Genotype V, but 5.2% distance to viruses from Genotype III. Based on previous estimates that classify DENV as distinct genotypes when the genomes differ by around 6%–8% at the nucleotide level [2,38], one might suggest that these viruses might be more appropriately classified as Genotype III. Finally, Dengue genotyping tool (available at https://www.genomedetective.com/app/typingtool/dengue/) was unable to classify the AF425625 at the genotype level, but AF425620 was classified as “*related but not part of DENV-1 Genotype III*”. We did not go further on the analysis of these two genomes but it is likely they represent either atypical Genotype III viruses or sylvatic strains, so we opted to include them as Genotype III in our tMRCA estimates.

## 5.0 Conclusion

In summary, this investigation has unveiled the presence of the two major lineages of DENV-1 Genotype V in Araraquara, SP and described two clade replacements within this Genotype during 2015-2022. These lineages exhibit a history of propagation across various regions of the country before emerging in the municipality. The implications are suggestive that these lineages might have influenced the divergent case counts observed during each epidemic occurrence and played a critical role on the maintenance of DENV-1 circulation. These results emphasize the importance of genomic surveillance as an additional tool in understanding the relationship between viral lineages and the magnitude of outbreaks. Furthermore, our findings have yielded novel insights into the origins of the DENV-1 genotype V – a strain of utmost significance within the Brazilian context.

## 6.0 Acknowledgements

This work was supported by FAPESP grant 2019/03859-9; by Medical Research Council and CADDE partnership award (MR/S0195/1) and FAPESP grant 2018/14389-0.

